# Persistent enhancement of intrinsic neuronal excitability induced by transient divalent cation depletion

**DOI:** 10.64898/2026.01.14.699613

**Authors:** Konstantina Mylonaki, Salvatore Incontro, Michaël Russier, Dominique Debanne

**Affiliations:** Aix-Marseille Université, INSERM, CNRS, LAI, Marseille, France; Vanderbilt University, Nashville, USA

**Keywords:** Hippocampus, LTP-IE, neuronal excitability, plasticity, heterosynaptic depression, CaSR, CaMKII, IP3 receptor

## Abstract

Fluctuations in external calcium concentration [Ca^2+^]_e_ occur during the wake and sleep cycle and during intense neuronal activity. However, the incidence of these fluctuations on neuronal excitability is not precisely known. We show here that reducing divalent cation (Ca^2+^ and Mg^2+^) concentrations from 1.3/0.8 to 0.6/0.4 mM during 15-30 minutes induces long-term potentiation of intrinsic excitability (LTP-IE) in CA1 pyramidal neurons. LTP-IE induced by low divalent cations is associated with a hyperpolarization of the action potential threshold and constitutes a positive feed-back of brain activity. This plasticity requires Ca^2+^-sensing receptor (CaSR), IP3 receptor (IP3R) and calcium-calmodulin kinase II (CaMKII). In fact, LTP-IE was occluded in the presence of the calcilytic NPS-2143 and absent in CRISPR CaSR neurons. In addition, inhibiting IP3R with 2-APB and CaMKII with kin considerably reduced LTP-IE magnitude. LTP-IE and synaptic potentiation (LTP) induced by spike-timing-dependent plasticity (STDP) protocol were also found to depend on CaSR as they were totally absent in CRISPR CaSR neurons. Spontaneous excitatory synaptic activity was found to be reduced by ∼35% following LTP induced by STDP. Importantly, this drop of spontaneous activity was not observed in CRISPR CaSR neurons. Taken together, these results show that CaSR plays a critical role in LTP-IE induced by low [Ca^2+^]_e_ and [Mg^2+^]_e_ and as well as in LTP of synaptic transmission and intrinsic excitability induced by STDP.

## Introduction

External calcium concentration ([Ca^2+^]_e_) in the brain fluctuates between 1.2 and 0.8 mM across the sleep/wake cycle (Ding *et al*., 2016) and may fall to 0.1 mM during intense neural activity such as that observed during epilepsy. Drop of [Ca^2+^]_e_ enhances excitability due to the hyperpolarization of the action potential (AP) threshold and is mediated by calcium sensing receptor (CaSR) and Nav1.2 channels (Mylonaki *et al*., 2025).

External magnesium concentration [Mg^2+^]_e_ also varies during the sleep/wake cycle from ∼0.70 mM during wakefulness to ∼0.83 mM during sleep, reaching up to 1.14 mM during anesthesia (Ding *et al*., 2016). External magnesium plays fundamental role in brain physiology as it blocks NMDA receptors at hyperpolarized potentials (Ascher & Nowak, 2009) and facilitates GABA_A_ receptor activity (Möykkynen *et al*., 2001). Neuronal activity increases intracellular Mg^2+^ concentration (Yamanaka *et al*., 2015) while reduced [Mg^2+^]_e_ levels are observed in epileptic patients (Kirkland *et al*., 2018). Raising [Mg^2+^]_e_ has been shown to elevate AP threshold and reduce intrinsic excitability (Frankenhaeuser & Meves, 1958; Dribben *et al*., 2010).

Physiological calcium levels deeply modify synaptic plasticity rules induced by spike-timing (Inglebert *et al*., 2020; Chindemi *et al*., 2022). However, whether reducing [Ca^2+^]_e_ as observed during intense neuronal activity induces *per se* synaptic and intrinsic changes remains unknown.

In this study, we report that lowering divalent cation concentrations from physiological levels observed at rest (i.e., 1.3 / 0.8 mM) to levels observed during intense neuronal activity (i.e., 0.6 / 0.4 mM) induces long-term potentiation of intrinsic excitability (LTP-IE) but no synaptic changes in CA1 pyramidal neurons. LTP-IE is absent in the presence of the calcilytic NPS-2143 or in neurons transfected with CRISPR CaSR. In addition, synaptic and intrinsic potentiation induced by spike-timing-dependent protocol (STDP) are not observed in CRISPR CaSR transfected neurons. We also report a reduction in spontaneous excitatory synaptic activity in control neurons that is absent in CRISPR CaSR neurons, thus pointing out the critical role of CaSR in plasticity.

## Results

### Reducing divalent cations induces LTP-IE but not LTP

CA1 pyramidal neurons maintained in organotypic slice cultures were recorded in whole-cell configuration. To check the incidence of low divalent cations on synaptic transmission and intrinsic excitability, excitatory postsynaptic potentials (EPSPs) and action potentials (APs) were evoked at low frequency (0.1 Hz) in physiological [Ca^2+^]_e_ and [Mg^2+^]_e_ (1.3 / 0.8 mM). After obtaining a stable baseline for both parameters, [Ca^2+^]_e_ and [Mg^2+^]_e_ were reduced to 0.6/0.4 mM during 15 minutes (**Figure 1A**). The test of intrinsic excitability and synaptic transmission was suppressed during this period (**Figure 1A** and **1B**). Then, the control solution was applied again and the test of intrinsic excitability was restored. A clear enhancement of the number of AP was observed that persisted during ∼30 minutes (152 ± 14%, n = 18; **Figure 1A**). In contrast, no change was observed on synaptic transmission (**Figure 1B**) indicating that reducing divalent cations does not induce LTP. LTP-IE was associated with a significant hyperpolarization of the AP threshold (from −49.5 ± 0.4 to −50.8 ± 0.4 mV; **Figure 1C**). No change in input resistance was observed (114 ± 5 vs. 114 ± 6 MΩ, n = 18, 30 minutes after bath application of low divalent cations; **Supplementary Figure 1**). The absence of intrinsic stimulation had no effect on the induction of LTP-IE as in parallel experiments, LTP-IE is still induced when the stimulation is kept throughout the whole experiment (**Supplementary Figure 2**).

**Figure 1.**
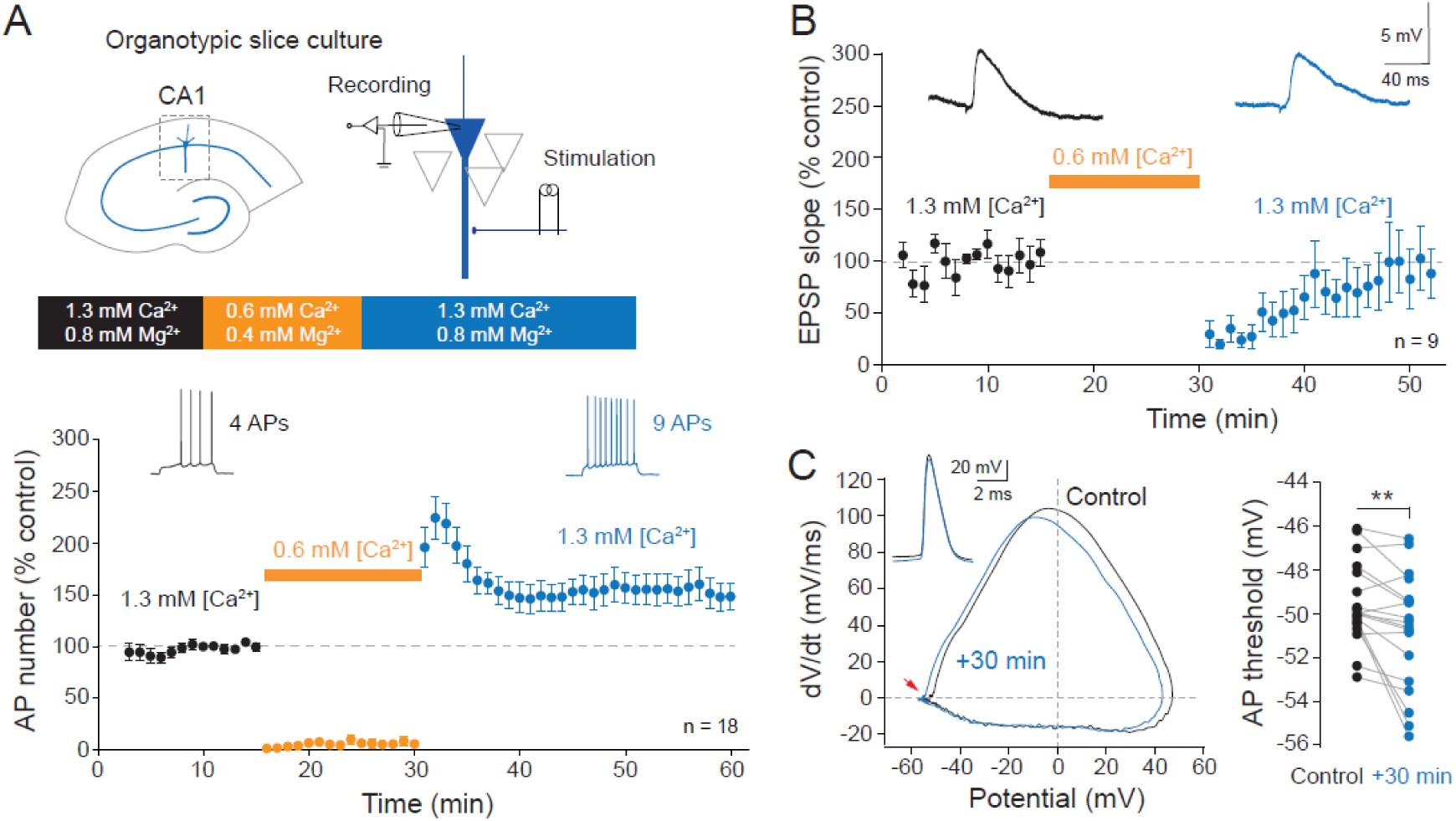
Low divalent cations induce LTP-IE but not LTP. A. Top, recording configuration and protocol for LTP-IE induction by transient reduction of divalent cations. Bottom, time-course of LTP-IE induced in control neurons. B. Lack of synaptic change induced by low divalent cations. C. Action potential threshold is persistently hyperpolarized following application of low divalent cations. **, Wilcoxon test, p<0.01.

In a second step, we tested whether the input-output function was altered by low divalent cations in a different set of experiment. The input-output curves was leftward shifted in the presence of low divalent cations and remained shifted when control levels of [Ca^2+^]_e_ and [Mg^2+^]_e_ were restored (**Figure 2A**). The rheobase remained also reduced in control solution (**Figure 2B**), confirming that excitability is altered on the long-term. The AP threshold was also found to be hyperpolarized after low divalent ion application (from −46.8 ± 0.5 to - 49.5 ± 0.8 mV; **Figure 2C**). The reduction of [Ca^2+^]_e_ alone from 1.3 mM to 0.6 mM in constant [Mg^2+^]_e_ (0.8 mM) was sufficient to induce a similar leftward shift of the input-output curve and a similar persistent reduction of the rheobase (**Supplementary Figure 3**). We conclude that reduction of [Ca^2+^]_e_ alone is sufficient to produce a persistent shift in intrinsic excitability.

**Figure 2.**
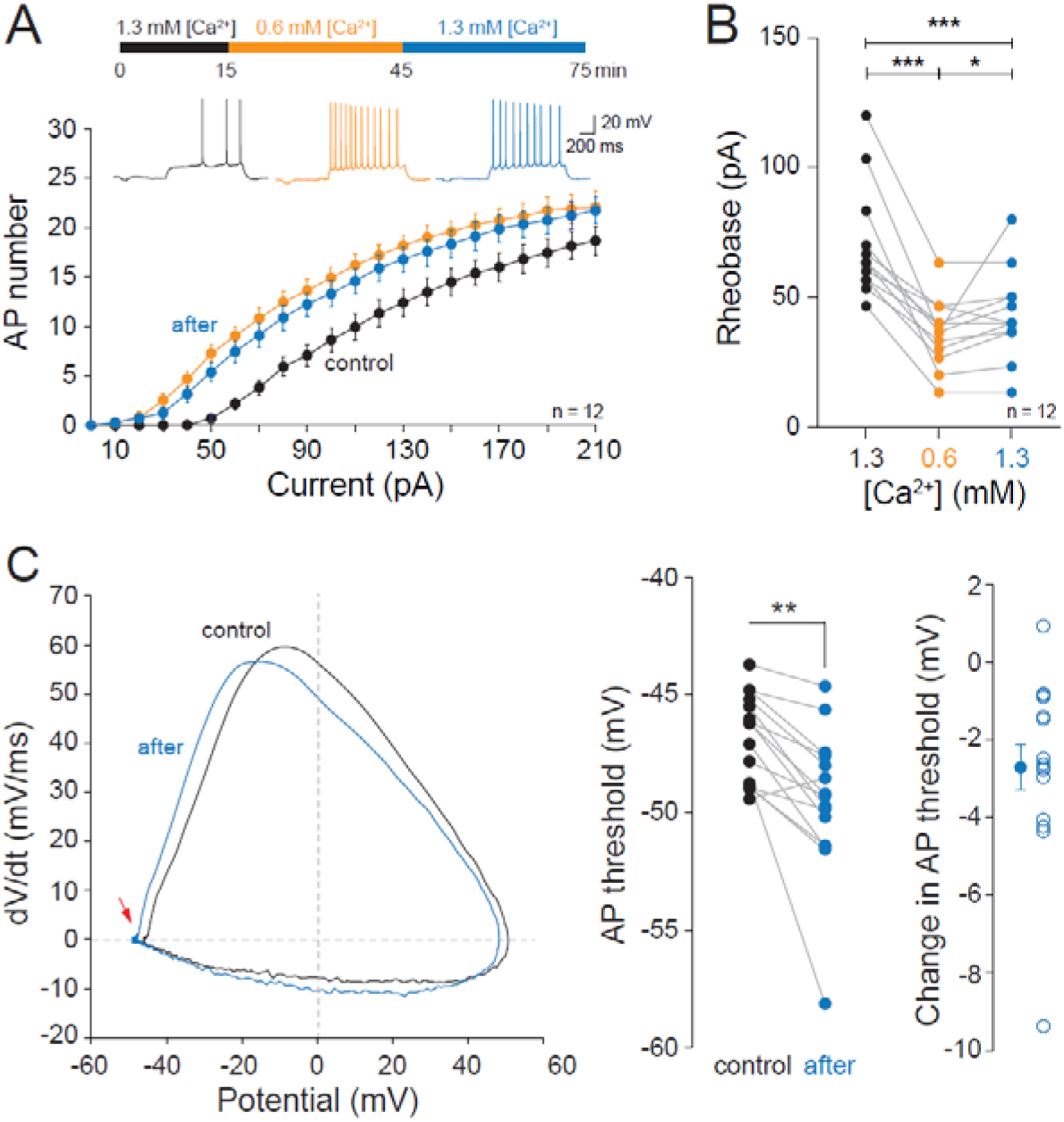
Input-output curves and rheobase analysis. A. Input-output curves in control (1.3/0.8 mM), low divalent ions (0.6/0.4 mM) and back in the control medium (after). Note the leftward shift in the curve after low divalent ions B. Rheobase change following low divalent ions. Note that the rheobase stay stable from 0.6 to 1.3 mM. ns, Wilcoxon test, p > 0.1. ***, Wilcoxon test, p<0.001. C. Hyperpolarization of the AP threshold. Left, phase plot of the AP before and after LTP-IE induction. Right, group data. **, Wilcoxon test, p<0.01.

Taken together, these results indicate that reduction of [Ca^2+^]_e_ induces LTP-IE in CA1 pyramidal neurons due to AP hyperpolarization.

### LTP-IE induced by low divalent cations requires CaSR

We next looked whether LTP-IE depended on CaSR. In the presence of the calcilytic agent NPS-2143, LTP-IE induced by transient low divalent depletion was totally abolished (**Figure 3A** and **Figure 3B**). Furthermore, in neurons transfected with CRISPR / CaSR (Mylonaki *et al*., 2025), no persistent shift in the input-output curves were observed nor persistent reduction in the rheobase (**Figure 3C** and **Figure 3D**). Altogether, these results indicate that CaSR activity is critical for LTP-IE induced by low divalent cations.

**Figure 3.**
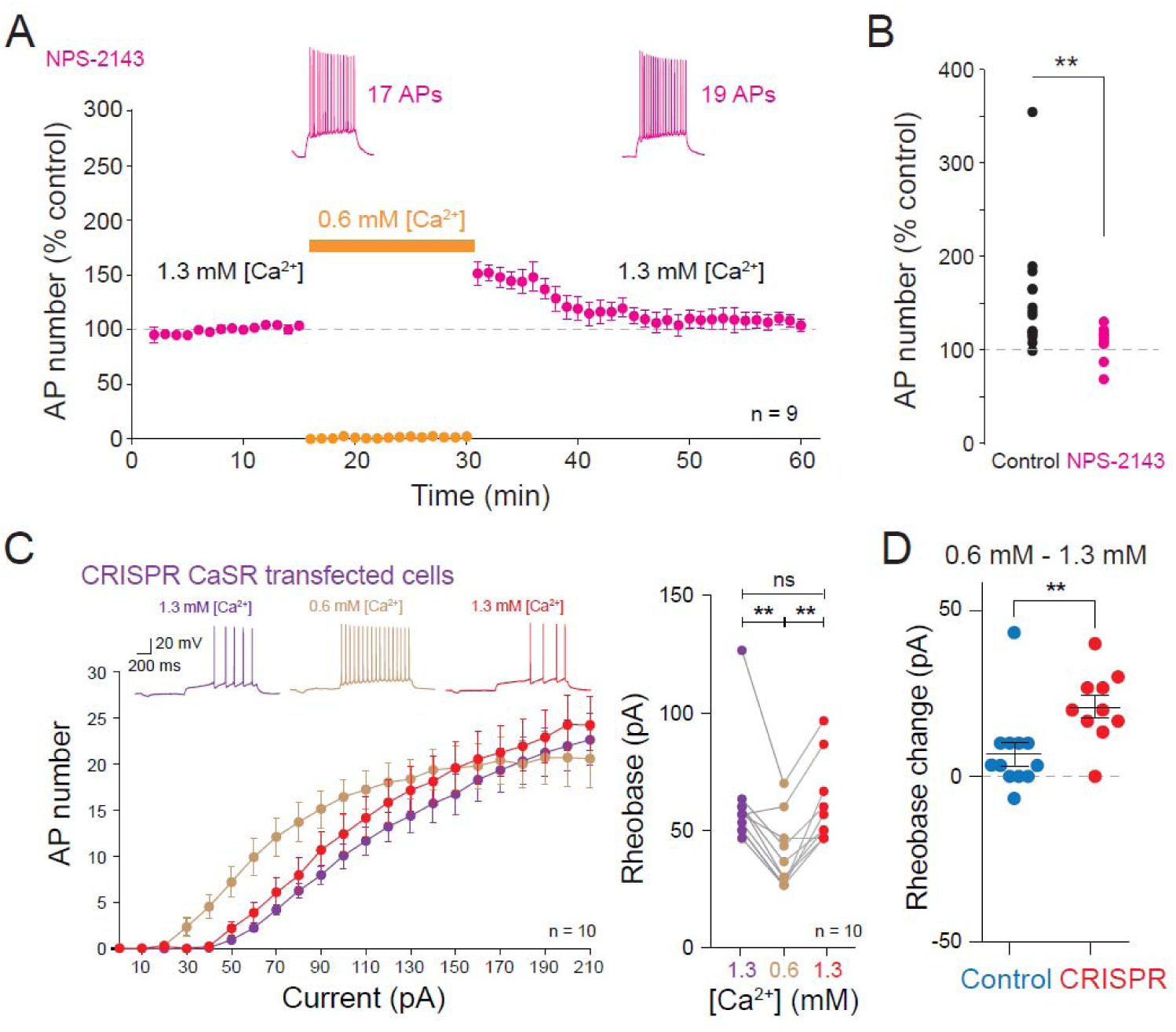
LTP-IE depends on CaSR activation. A. NPS-2143 (10 µM) occludes LTP-IE induced by low divalent cations. B. Comparison with control (i.e., no drug). **, Mann-Whitney U-test, p<0.01. C. No change in rheobase in neurons transfected with CRISPR CaSR. Left, input-output curves in control (1.3 mM calcium and 0.8 mM magnesium, violet data points), 0.6 mM calcium and 0.4 mM magnesium (light brown) and back to the control solution (1.3 mM calcium and 0.8 mM magnesium; red data points). Right, analysis of the rheobase. **, Wilcoxon test, p<0.01. D. Difference in the rheobase from 0.6 to 1.3 mM in control (blue points) and CRISPR CaSR (red data points). **, Mann-Whitney U-test, p<0.01.

### LTP-IE induced by low divalent cations requires IP3 receptor and CaMKII activation

CaSR activates the phospholipase C (PLC) that in turn mobilizes calcium from internal stores through activation of IP3 receptors. We therefore tested the role of IP3R using a specific inhibitor, 2-APB. In the presence of 2-APB, LTP-IE was totally occluded (**Figure 4A** and **Figure 4B**). Calcium released by internal calcium stores may in turn activates CaMKII. We next tested the effect of CaMKII inhibition on LTP-IE induction. CaMKII inhibition with KN93 largely occluded LTP-IE (**Figure 4C**) indicating that CaMKII is involved in the persistent increase in intrinsic excitability. Thus, a possible scheme would be that CaSR activates the production of IP3 that opens calcium stores that activates CaMKII to induce LTP-IE (**Figure 4D**).

**Figure 4.**
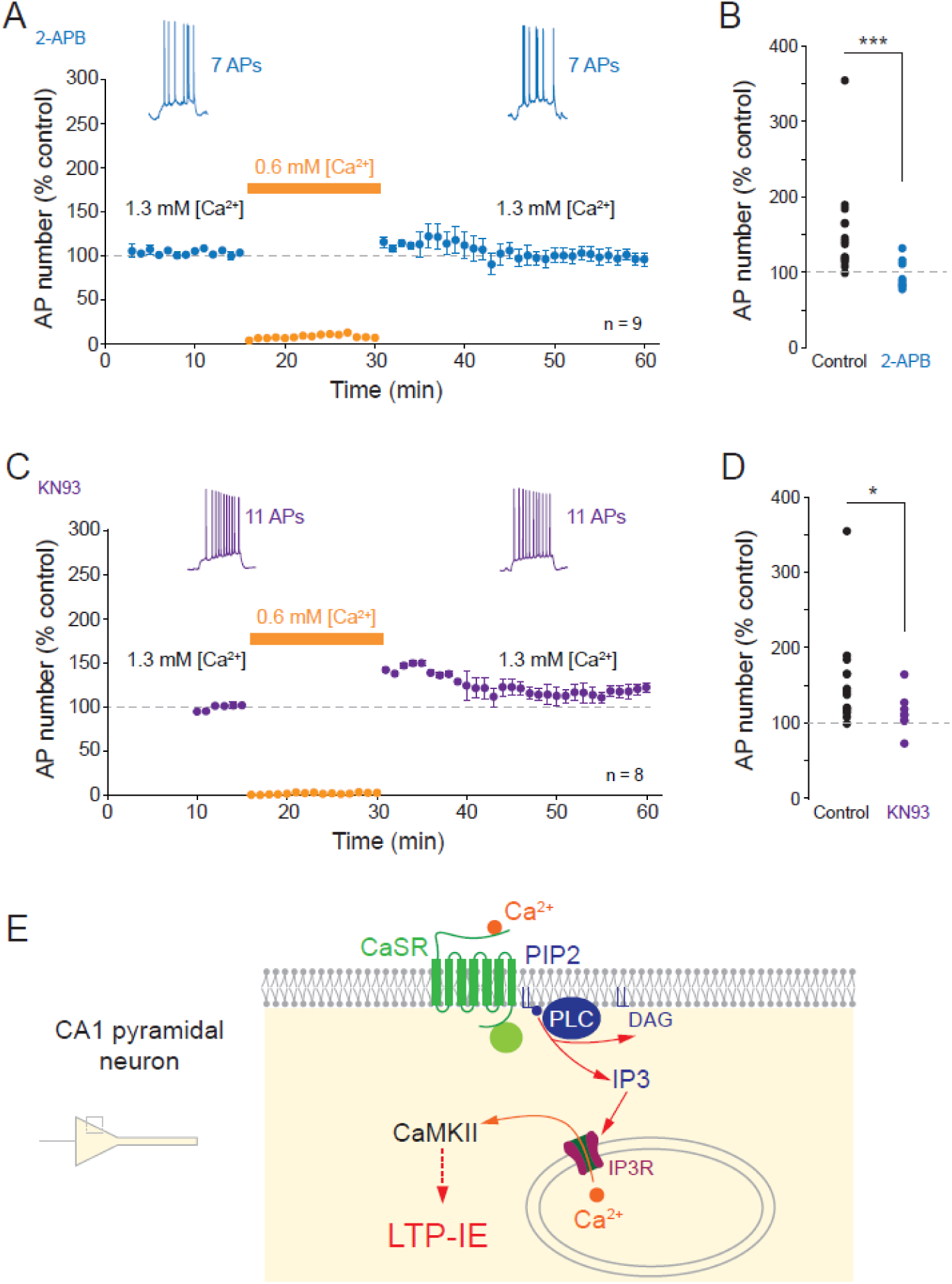
LTP-IE depends on IP3R and CaMKII. A. LTP-IE induced by low divalent ions is blocked in the presence of the IP3R inhibiter 2-APB (15 µM). B. Comparison with the control (i.e., no drug). ***, Mann Whitney U-test, p<0.001. C. LTP-IE induced by low divalent ions is prevented in the presence of the CaMKII inhibiter, KN93 (10 µM). D. Comparison with the control (i.e., no drug). *, Mann-Whitney U-test, p<0.05. E. Possible scenario of LTP-IE induction by low divalent ions involving CaSR, PIP2 hydrolysis, IP3 receptor activation and CaMKII.

### LTP and LTP-IE induced by STDP requires CaSR

We next looked whether CaSR is required in LTP-IE induced by spike-timing-dependent plasticity protocol (STDP) in which single EPSP is paired with a single postsynaptic AP with a delay of +10 ms at a frequency of 10 Hz. In control neurons, LTP and LTP-IE are induced (**Figure 5A** and **Figure 5B**). In contrast, in CRISPR CaSR neurons both forms of plasticity were absent (**Figure 5C** and **Figure 5D**) and the levels of synaptic and intrinsic potentiation were found to be significantly reduced compared to control neurons in CRISPR CaSR neurons (**Figure 5E** and **Figure 5F**). Excitability was stable when no STDP protocol was applied (**Supplementary Figure 4**). Taken together, these results confirm the importance of CaSR in functional plasticity.

**Figure 5.**
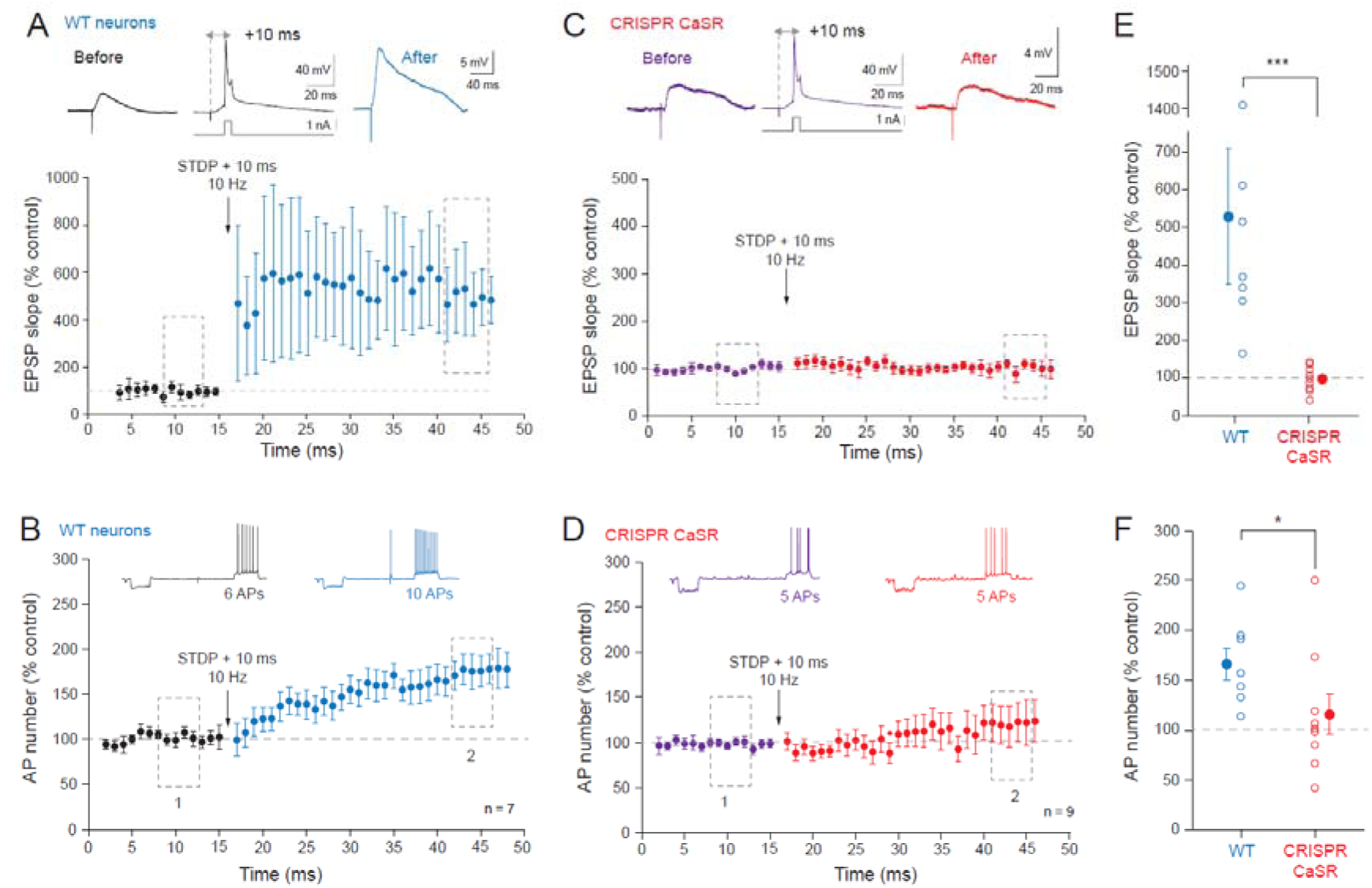
LTP and LTP-IE induced by STDP are prevented in CRISPR CaSR neurons. A. LTP induction in wild-type (WT) neurons. Top representative EPSP before and after pairing with a delay of +10 ms between the EPSP and the postsynaptic AP at 10 Hz. B. LTP-IE induced in parallel with LTP in WT neurons. C. No LTP is induced in CRISPR CaSR neurons. D. No LTP-IE is induced in CRISPR CaSR neurons. E. Comparison of LTP induced in WT and CRISPR CaSR neurons. ***, Mann-Whitney U-test, p<0.001. F. Comparison of LTP-IE induced in WT and in CRISPR CaSR neurons. *, Mann-Whitney U-test, p<0.05.

### Long-lasting reduction of spontaneous activity requires CaSR

In physiological calcium, spontaneous excitatory synaptic activity measured in time window before synaptic stimulation (**Supplementary Figure 5**) is relatively high (mean frequency: ∼6 Hz; **Figure 6A**) and episodes of oscillation at θ and γ frequency were observed (**Supplementary Figure 6**). The amplitude of spontaneous EPSPs was on average ∼1.4 mV, ranging from 0.1 to 11.4 mV (**Figure 6B**). A marked reduction of this spontaneous activity (by ∼35%) was observed following induction of LTP with the STDP protocol (**Figure 6A**). This reduction lasted at least 40 minutes post STDP. Interestingly, no difference was observed in the amplitude of spontaneous EPSPs before and after LTP induction (**Figure 6B**), indicating that the reduction in spontaneous event frequency does not correspond to a reduction in the release probability but rather to the reduction in the firing frequency of presynaptic neurons that were not involved in synaptic potentiation (**Figure 6C**). We conclude that CaSR plays a critical role in the long-lasting modulation of circuit activity.

**Figure 6.**
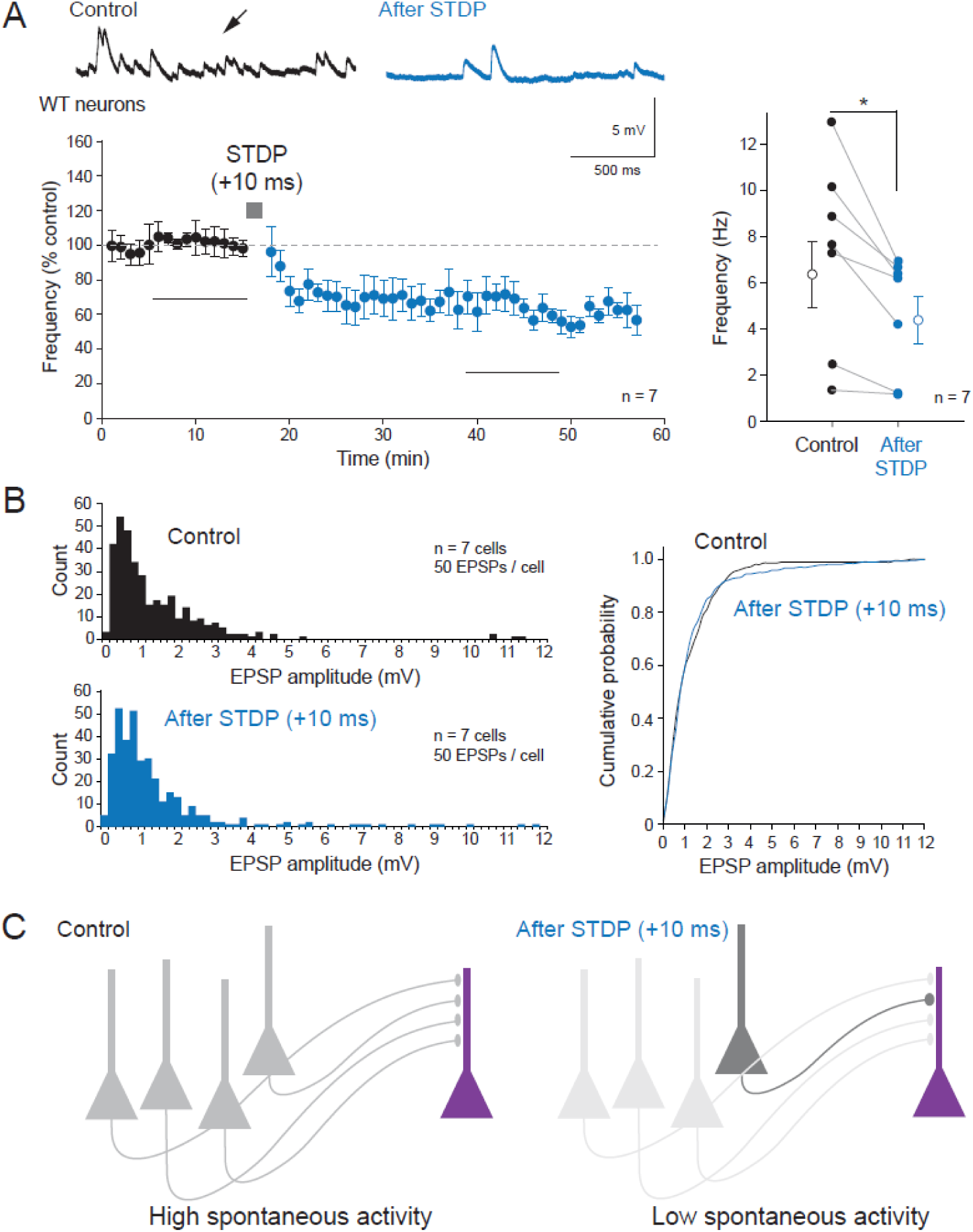
Reduction of the spontaneous activity after LTP induction. A. Left, time-course of the spontaneous activity recorded in control neurons before and after LTP induced by STDP. Right, quantification made during the last 10 minutes in control and after LTP. *, Wilcoxon test, p<0.05. B. Lack of EPSP amplitude change. Left, histograms of EPSP amplitude before (top) and 20-30 minutes after (bottom) induction of LTP. Right, comparison of the cumulative distributions of EPSPs before and after LTP. C. Possible scenario accounting for the reduction of spontaneous activity. Before LTP, all presynaptic cells fire spontaneously accounting for spontaneous activity. When LTP is induced at the synapse formed by one of the input cells, this cell expresses an enhanced excitability symbolized by dark grey (Ganguly *et al*., 2000; Li *et al*., 2004) while the other presynaptic cells would undergo a reduction in excitability (symbolized by light grey), thus accounting for the low spontaneous activity observed.

Next, we checked whether excitatory spontaneous activity was also reduced in CRISPR CaSR neurons. In contrast with WT neurons, no change in spontaneous activity was observed in neurons transfected with CRISPR CaSR (**Figure 7A**). A significant difference in the frequency change was found between WT and CRISPR CaSR neurons (**Figure 7B**). These data indicate that reduction of spontaneous activity is also controlled by CaSR.

**Figure 7.**
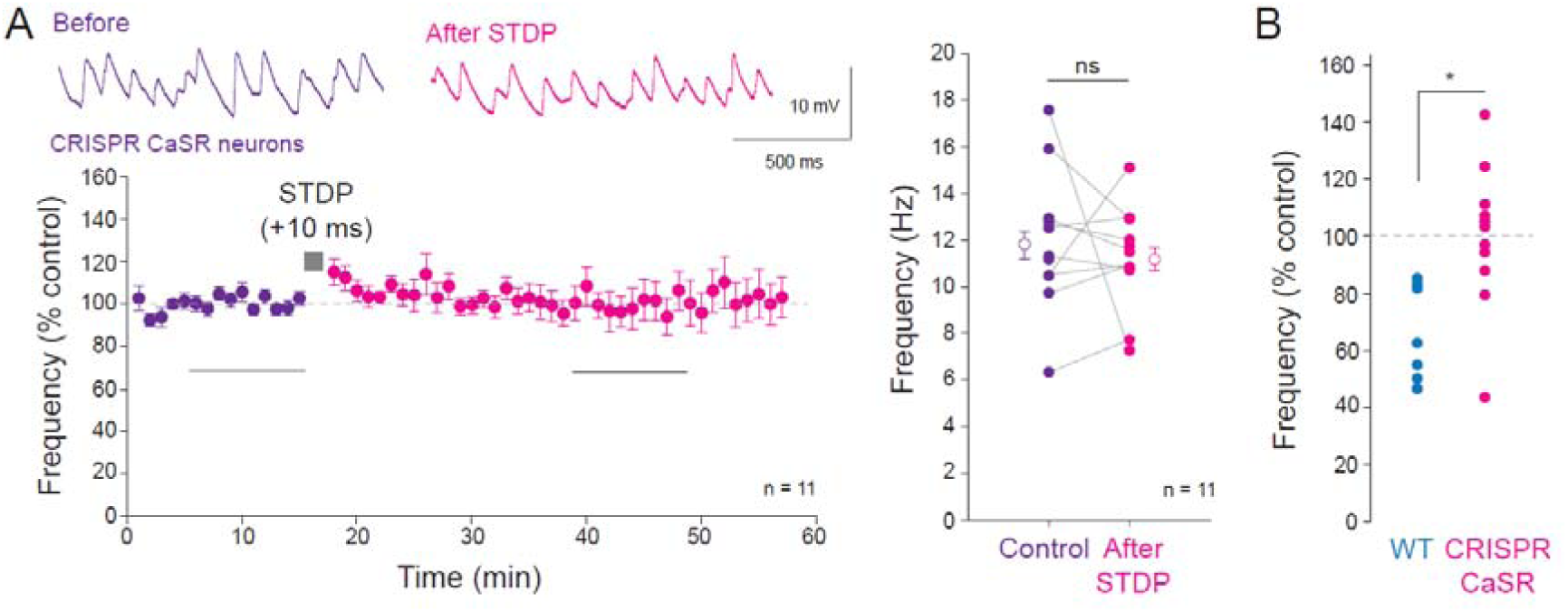
Spontaneous activity is stable following STDP induced in CRISPR CaSR neurons. A. Left, time course of the frequency of spontaneous excitatory synaptic activity before and after STDP (+10 ms). Right, group data. ns, Wilcoxon, p>0.1. B. Comparison of the normalized frequency of spontaneous activity in wild type (WT) and CRISPR CaSR neurons. *, Mann-Whitney U-test, p<0.05.

## Discussion

We show here that reducing divalent cation (i.e., Ca^2+^ and Mg^2+^) concentration from 1.3/0.8 to 0.6/0.4 induces long-lasting enhancement of intrinsic excitability (LTP-IE) in CA1 hippocampal neurons. This form of LTP-IE is prevented in the presence of the calcilytic NPS-2143 or in CRISPR CaSR neurons, indicating that CaSR plays a critical role in the induction of LTP-IE. In addition, LTP-IE is prevented by blocking IP3R and CaMKII indicating that it depends on internal calcium stores and CaMKII. Furthermore, we show that both LTP and LTP-IE induced by STDP (+10 ms) are absent in CRISPR CaSR neurons. Finally, the reduction in spontaneous excitatory synaptic activity is also prevented in CRISPR CaSR neurons, indicating that CaSR activation represents a critical step in this plasticity.

### External calcium and magnesium depletion

External calcium depletion can be produced by synaptic activity (Borst & Sakmann, 1999; Rusakov & Fine, 2003; Cohen & Fields, 2004) or by epileptiform activity (Pumain *et al*., 1983; Raimondo *et al*., 2015). The calcium depletion caused by synaptic activity lasts a few tens of milliseconds while the depletion caused epileptic activity is much longer (>2 s for postictal spikes). External magnesium depletion occurs during synaptic activity and epilepsy (Yamanaka *et al*., 2015; Kirkland *et al*., 2018).

In our experiments, the depletion of both calcium and magnesium to 0.6 and 0.4 mM is aimed to mimic the changes in external divalent cation concentration due to medium range neuronal activity observed during wakefulness (Ding *et al*., 2016). The role of magnesium drop *per se* is minor in LTP-IE induction since no difference in the rheobase shift was observed when calcium concentration was dropped alone versus calcium and magnesium were both reduced. The acute effect of magnesium reduction in physiological conditions was also found to be negligeable (Mylonaki *et al*., 2025).

### LTP-IE induced by transient drop of divalent cation concentration

Long-lasting plasticity of intrinsic neuronal excitability can be classically induced by synaptic stimulation (Sourdet *et al*., 2003; Xu *et al*., 2005; Campanac *et al*., 2013; Incontro *et al*., 2021; Sammari *et al*., 2022; Duménieu *et al*., 2025) or by spiking activity (Cudmore & Turrigiano, 2004; Nataraj *et al*., 2010; Duménieu *et al*., 2025). Here, we show that LTP-IE can also be induced by transiently lowering external calcium concentration. The induction mechanisms involve activation of CaSR and of IP3R to mobilize calcium from internal stores and activation of CaMKII. The expression mechanisms have not been precisely explored here but the AP threshold which depends on Nav and Kv1 channels was found to be hyperpolarized. Depletion of calcium stores has been shown to induce lasting regulation of h-channels in CA1 pyramidal neurons (Narayanan *et al*., 2010). While LTP-IE is generally induced in parallel with LTP (Debanne *et al*., 2019), no change in synaptic transmission was observed following depletion of divalent cations. The mechanism underlying this differential effect is unknown but it may result from insufficient CaMKII activation to induce synaptic plasticity.

Modulation of ion channels has been demonstrated in the context of epilepsy (Beck & Yaari, 2008; Debanne *et al*., 2024). Some changes are anti-epileptic (homeostatic) while others are pro-epileptic (Hebbian). The mechanism reported here corresponds to a broad Hebbian mechanism as it could enhance neuronal activity following a period of low calcium due to active circuits.

### Long-lasting reduction of spontaneous activity during LTP

We report here that LTP induced by positive pairing (+10 ms) induces a marked reduction of spontaneous activity in the recorded neuron. In contrast with the great majority of previous studies, the circuit is intact (i.e., no cut between CA3 and CA1), the synaptic inhibition is preserved and the external calcium concentration is physiological (1.3 mM). At first glance, this reduction could correspond to the well-known heterosynaptic depression reported by many authors at non stimulated inputs when LTP is induced at one input (Jenks et al., 2021). However, heterosynaptic depression is mostly presynaptic (Huang *et al*., 2008; Smith *et al*., 2020) and the reduction in spontaneous activity observed in our experiments does not seem to corresponds to a reduction in miniature events because i) the amplitude distribution of spontaneous EPSPs is similar to the distribution of unitary synapses made by CA3 and CA1 pyramidal neurons in organotypic slice cultures (Debanne *et al*., 1995), and ii) the EPSP amplitude distribution remained unchanged following LTP induction. This argues for a reduction in spontaneous firing of presynaptic neurons that did not participate to the synaptic potentiation. Importantly, LTP induced by STDP is accompanied by a long-lasting increase in presynaptic excitability (Ganguly *et al*., 2000; Li *et al*., 2004). Whether other presynaptic neurons impinging on the postsynaptic cell display a reduction of intrinsic excitability is not yet known. A possible scenario depicted in **Figure 6** envisages that presynaptic neurons that are not involved in the potentiated input would undergo a reduction of intrinsic neuronal excitability, thus accounting for the reduced spontaneous activity. Supporting this view, the fiber volley corresponding to the presynaptic APs recorded extracellularly is depressed on the non-stimulated pathway that undergo heterosynaptic plasticity (Yasuda *et al*., 2008). This presynaptic depression depends on potassium channels and thus can be seen as a reduction in intrinsic excitability of non-potentiated inputs. This hypothesis implies the existence of a retrograde messenger produced by the postsynaptic neuron or by nearby astrocytes and acting selectively on non-activated neurons. Further studies will be required to identify the precise mechanisms.

### Calcium sensing receptor and learning and memory

Although CaSR plays a critical role in decision making in vertebrate (Shoenhard *et al*., 2022), the role of CaSR in learning and memory remains largely unknown. The β-amyloid peptide is known to activate CaSR (Conley *et al*., 2009) and inhibition of CaSR improves spatial memory deficits in mouse models of Alzheimer disease (Feng *et al*., 2020). CaSR is involved in the regulation of synaptic transmission (Phillips *et al*., 2008). However, the mechanisms linking synaptic / intrinsic plasticity and CaSR was still lacking. We believe that the present study fills this gap by showing the critical role of CaSR i) on intrinsic excitability induced by low divalent cations and ii) on synaptic and intrinsic plasticity induced by STDP.

## Materials and methods

### Organotypic slice cultures of rat hippocampus

All experiments were carried out according to the European and Institutional guidelines for the care and use of laboratory animals (Council Directive 86/609/EEC and French National Research Council) and approved by the local health authority (CE71, D13055-08, Préfecture des Bouches-du-Rhône). Slices cultures were prepared as described previously (Debanne *et al*., 2008). In brief, young Wistar rats (P7–P10) were deeply anesthetized with isoflurane and killed by decapitation, the brain was removed, and each hippocampus was dissected in a sodium-free solution (in mM): Sucrose, 283; NaHCO_₃_, 26; D-Glucose, 10; KCl, 1.3; CaCl_₂_, 1; MgCl_₂_, 10; Kynurate, 200 mM continuously equilibrated with 95% O_2_ – 5% CO_2_. Hippocampal slices (350 μm) were obtained using a Vibratome (Leica, VT1200S). They were placed on 20-mm latex membranes (Millicell) inserted into 35-mm Petri dishes containing 1 mL of culture medium and maintained for up to 18 d in an incubator at 35°C, 95% O_2_–5% CO_2_. The culture medium contained 25 ml MEM, 1.25 ml HBSS, 12.5 ml horse serum, 0.5 ml penicillin/streptomycin, 0.8 ml glucose (1 M), 0.1 ml ascorbic acid (1 mg/ml), 0.4 ml Hepes (1 M), 0.5 ml B27, and 8.95 ml sterile H_2_O.

### Electrophysiology

After 10-14 DIV, whole-cell recordings were obtained from electroporated CA1 neurons, which were identified by GFP expression. The perfusion solution contained (in mM): 125 NaCl, 2.5 KCl, 0.8 NaH_2_PO_4_, 26 NaHCO_3_, 10 glucose, 0.8 MgCl_2_ and 1.3 CaCl_2_ continuously equilibrated with 95% O_2_ – 5% CO_2_. CA1 pyramidal hippocampal neurons were identified by the location of their soma, their morphology and their electrophysiological profile. Patch pipettes (7–9 MV) were filled with the internal solution containing (in mM): 120 K-gluconate, 20 KCl, 10 HEPES, 0 EGTA, 2 MgCl_2_, 2 Na_2_ATP, and 0.3 NaGTP (pH 7.4). All recordings were made at 29°C in a temperature-controlled recording chamber (Luigs & Neumann GmbH) with oxygenated ACSF. Slice cultures were kept intact (i.e., without surgical removal of area CA3) and recordings from CA1 pyramidal neurons were made in the presence of intact inhibition (without PiTx) Neurons were recorded in current clamp with a Multiclamp 700B Amplifier (Molecular Devices). Excitability was measured by delivering a range of long (1 s) depolarizing current pulses (10– 290 pA, by increments of 10 pA) and counting the number of action potentials. In most experiments, the ratio of CaCl_2_ and MgCl_2_ concentration was maintained constant, and [Ca^2+^]_e_ was set to either 1.3 mM ([Mg^2+^]_e_ = 0.8 mM), or 0.6 mM ([Mg^2+^]_e_ = 0.4 mM). In some experiments, the [Mg^2+^]_e_ was kept constant.

Input-output curves were determined for each neuron and three parameters were examined: the rheobase (the minimal current eliciting at least one action potential), the amplitude of the Action potential and the latency of the first spike (depolarizing time before the evoked spike under rheobase current eliciting only 1 spike). The current signals were low-pass filtered (10 kHz,), and acquisition was performed at 10 kHz with pClamp10 (Molecular Devices). Data were analyzed with ClampFit (Molecular Devices) and Igor (Wavemetrics). Spike thresholds were measured using phase plots (Yu *et al*., 2008; Fékété *et al*., 2021). Synaptic stimulation was evoked at low frequency (0.1 Hz) by a stimulating electrode positioned in the stratum radiatum (Inglebert *et al*., 2020). LTP and LTP-IE were induced with EPSPs paired with single postsynaptic AP (delay: +10 ms) at 10 Hz.

Spontaneous EPSPs were detected as fast upward voltage transitions during the time period of 1.8 sec before synaptic stimulation (**Supplementary Figure 5**). The minimal detection threshold was set to twice the recording noise value (∼0.1 mV).

### CRISPR/Cas9

The primers used to design the specific gRNA targets were: CaSR forward: (5′ to 3′) CACC G ctgctactccaaaatggat; CaSR reverse (3’ to 5’) AAAC atccattttggagtagcaag C. The gRNA sequences were ligated into pX458 to co-express the human codon-optimized Cas9 as previously described (Incontro *et al*., 2014). We used pX458 expressing gRNA targeting CaSR and FUGW expressing plasmid to aid identification of transfected cells. The human codon-optimized Cas9 and chimeric gRNA expression plasmid (pX458) as well as the lentiviral backbone plasmid (lentiCRISPR) both developed by the Zhang lab (Cong *et al*., 2013; Ran *et al*., 2013) were obtained from Addgene. To generate guide RNA (gRNA) plasmids, a pair of annealed oligos (20 bp) was ligated into the single gRNA scaffold of pX458 or lentiCRISPR. For all the experiments, we sub-cloned the two gRNAs including the following chimeric RNA into a pCAGGS-IRES-GFP plasmid, through PCR amplification and insertion into the plasmid using BstbI restriction sites. To enhance the identification of transfected neurons we co-expressed the pFUGW vector expressing only GFP with pCAGGS-IRES-GFP constructs.

### Single cell electroporation

Electroporation-mediated transfection (Rathenberg *et al*., 2003) was conducted in organotypic slice cultures of rat hippocampus at 4 d in vitro (DIV). Before electroporation, plasmid solutions were centrifuged at 10,000 g for 5 min to avoid obstruction of the micropipette. For single-cell electroporation (SCE), the microscope chamber was consisting of a sterile 35-mm Petri dish. The plasmid DNA constructs were diluted to a final concentration of 33 ng/ul in the internal solution containing (in mM): 120 K-gluconate, 20 KCl, 10 HEPES, 0 EGTA, 2 MgCl_2_, 2 Na_2_ATP, and 0.3 NaGTP (pH 7.4). Micropipettes (7–10 MΩ) were filled with this DNA preparation after filtration with a sterile Acrodisc Syringe Filter (0.2 mm in pore diameter). During the SCE procedure, slice culture (DIV–DIV4) was positioned in the 35-mm Petri dish and covered with prewarmed and filtered external solution containing (in mM): 125 NaCl, 26 NaHCO_3_, 3 CaCl_2_, 2.5 KCl, 2 MgCl_2_, 0.8 NaH_2_PO_4_, 0.6 Hepes and 10 D-glucose, equilibrated with 95% O_2_–5% CO_2_. The ground electrode and the microelectrode were connected to an isolated voltage stimulator (Axoporator 800A, Molecular Devices). Under visual guidance, the micropipette was positioned by a three-axis micromanipulator near the cell body of selected CA1 neurons. Pressure was controlled to have a loose-seal between the micropipette and the plasma membrane. When the resistance monitored reached 25–30 MΩ, we induced a train of −12 V pulses during 500 ms (pulse width: 0.5 ms, frequency: 50 Hz). Each organotypic slice culture underwent SCE procedure for 5–10 selected neurons during a limited time of 30 min and was then back transferred to the incubator.

### Pharmacology

NPS 2143 hydrochloride (2-Chloro-6-[(2R)-3-[[1,1-dimethyl-2-(2-naphthalenyl) ethyl] amino-2-hydroxypropoxy] benzonitrile hydrochloride), 2-Aminoethoxydiphenylborate, (2-Aminoethoxy) diphenylborane (2-APB), and (E)-N-(2-(((3-(4-chlorophenyl) allyl) (methyl)amino) methyl) phenyl)-N-(2-hydroxyethyl)-4-methoxybenzenesulfonamide (KN93) were purchased from Tocris.

### Statistics

Pooled data are presented as mean ± SEM and statistical analysis was performed using the Mann–Whitney *U*-test or Wilcoxon rank-signed test. Data were considered statistically significant for p < 0.05.

## Acknowledgments

We thank K. Milton and A. Venture for animal care and L. Fronzaroli-Molinieres, N. Boumedine-Guignon and M. Sangiardi for excellent technical assistance.

## Author contributions

K.M. designed research, performed research, analyzed the data, built the figures and wrote the paper. S.I. performed research and analyzed the data, M.R. analyzed the data. D.D. designed research, analyzed the data, built the figures, provided funding and wrote the paper.

## Data availability

All data needed to evaluate the conclusions in the paper are present in the manuscript and/or the Supplementary Materials.

---

**Supplementary Figure 1.**
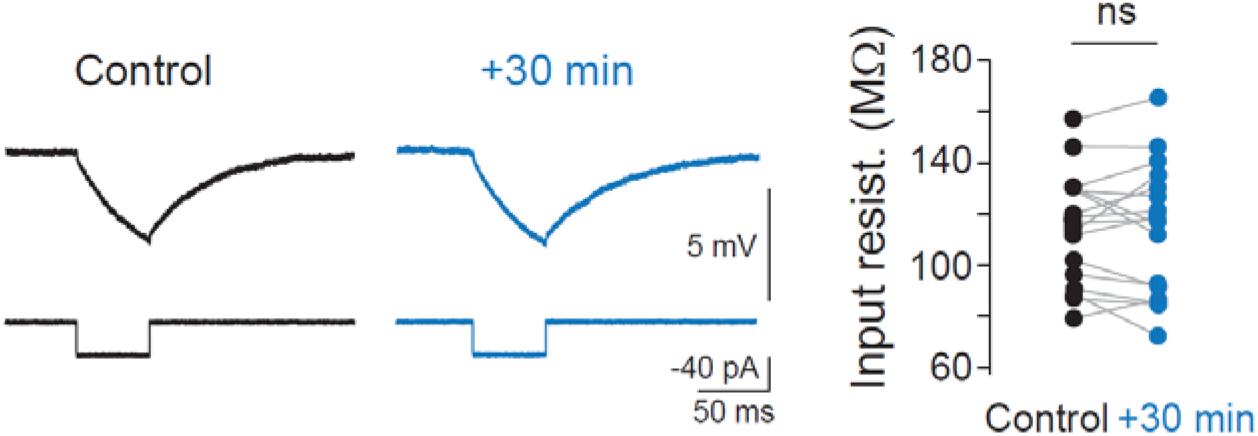
Input resistance remains unchanged following application of low divalent cations. Left, representative voltage traces before and 30 minutes after application of low divalent ions for 15 minutes. Right, quantitative data. ns, Wilcoxon test, p>0.1.

**Supplementary Figure 2.**
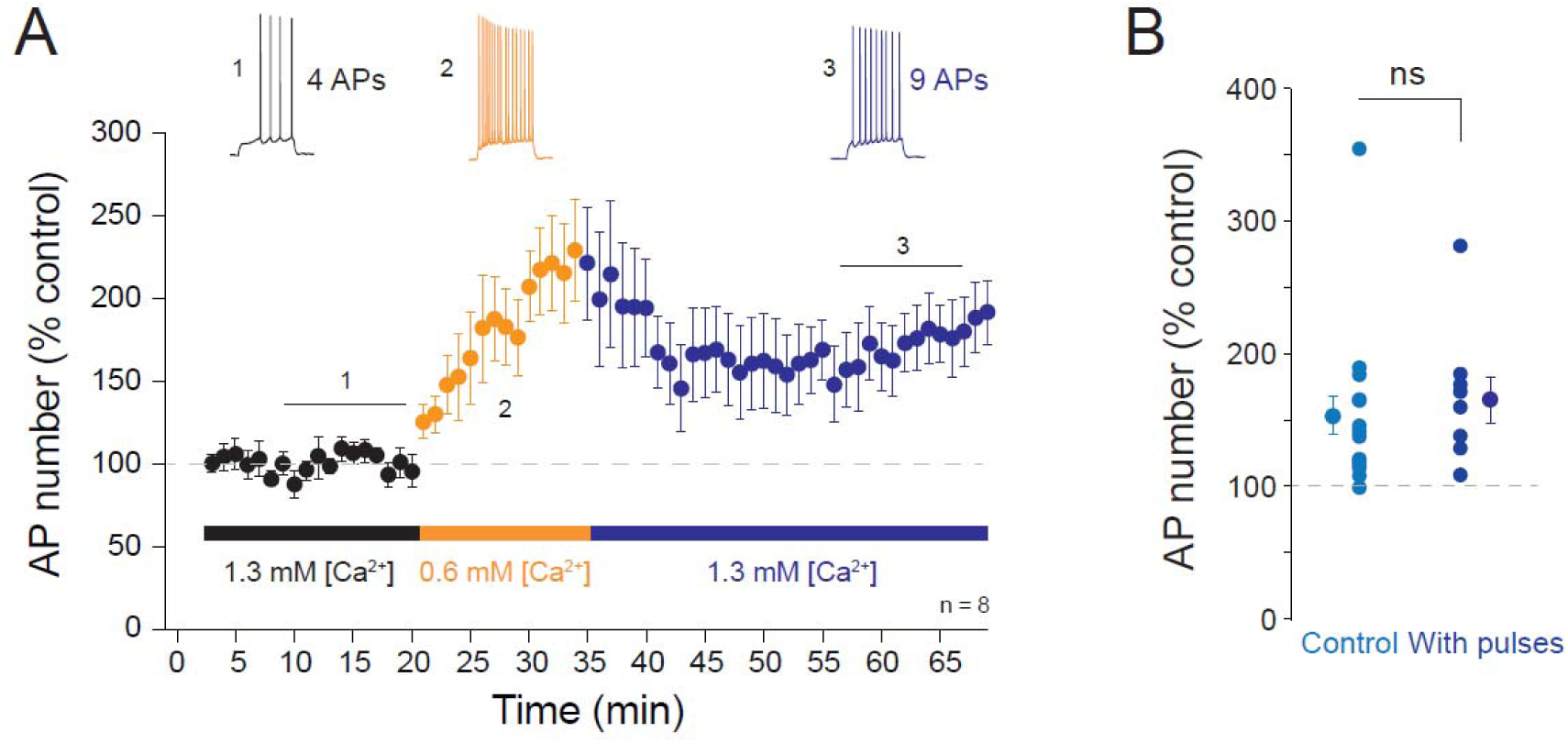
Role of pulses on LTP-IE induced by low divalent cations. A. Time-course of the AP number before and after application of low divalent cations (0.6/0.4 mM). B. Comparison of the results obtained with pulses with the control data (without pulses). ns, Mann-Whitney U-test p>0.1.

**Supplementary Figure 3.**
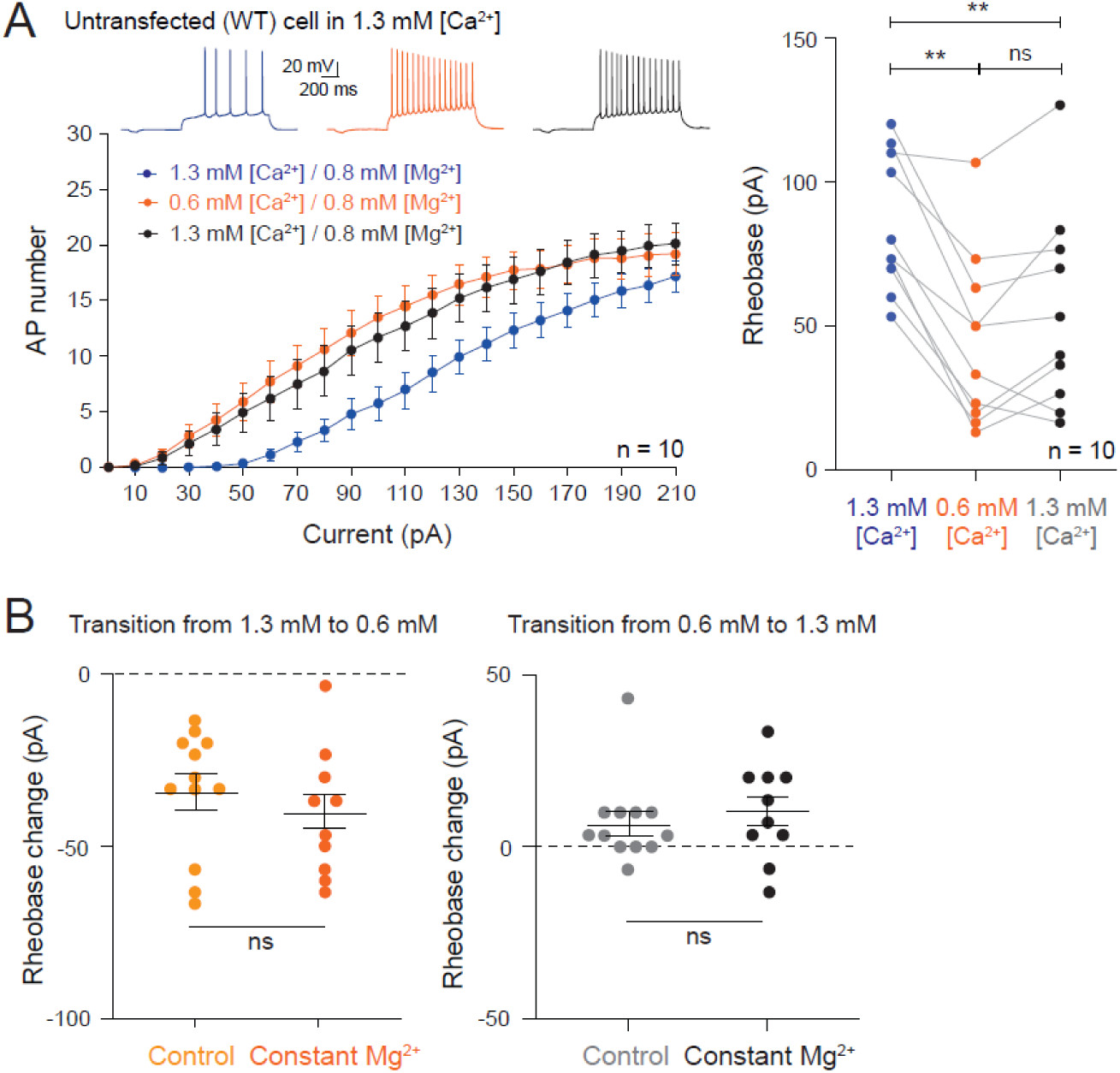
LTP-IE induced by low [Ca^2+^]_e_ in constant [Mg^2+^]_e_. A. Left, input-output curves remain left shifted following reduction of [Ca^2+^]_e_. Right, rheobase remains persistently reduced following low calcium. **, Wilcoxon test, p<0.01. ns, Wilcoxon test, p>0.05. B. Left, rheobase change during 0.6 mM [Ca^2+^]_e_. Right, rheobase change during the recovery in 1.3 mM [Ca^2+^]_e_. ns, Mann-Whitney U-test, p>0.1.

**Supplementary Figure 4.**
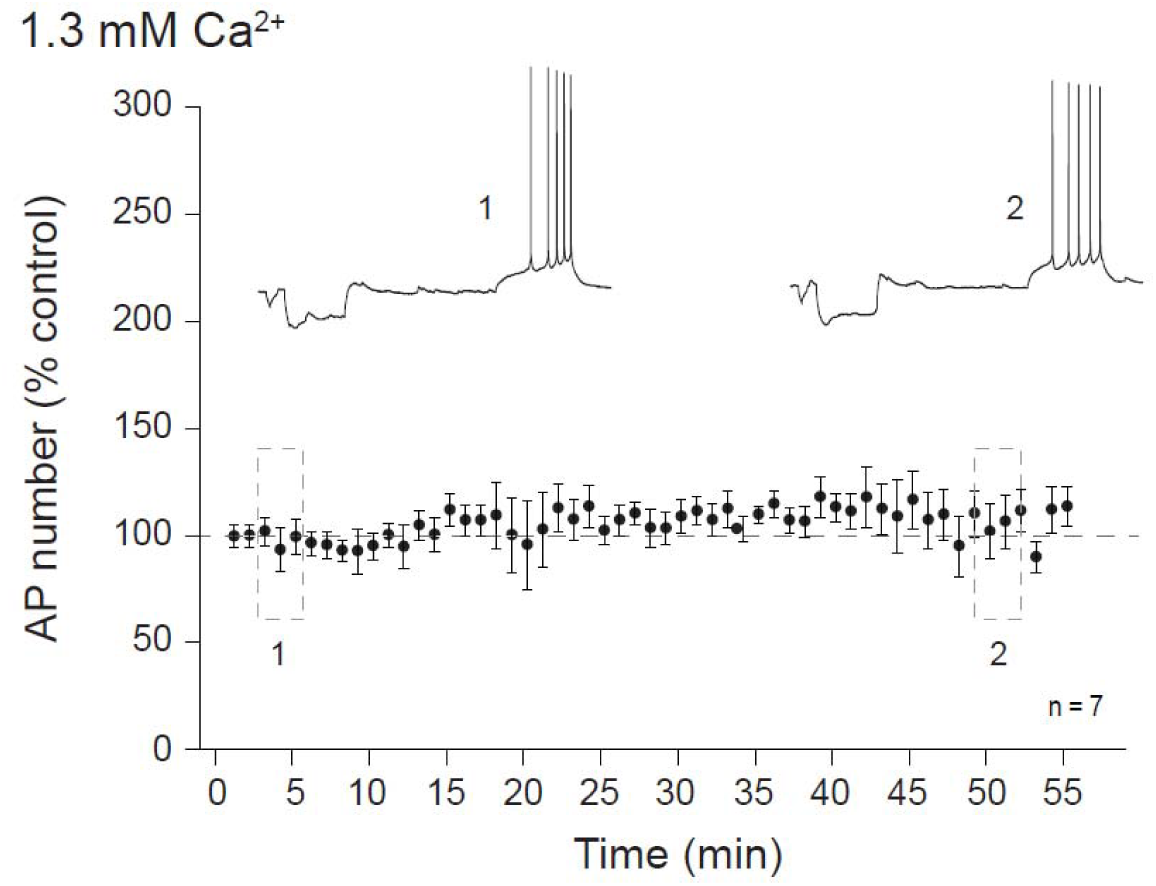
Temporal stability of intrinsic neuronal excitability. Temporal stability of the intrinsic neuronal excitability recorded in CA1 pyramidal neurons from organotypic slice cultures.

**Supplementary Figure 5.**
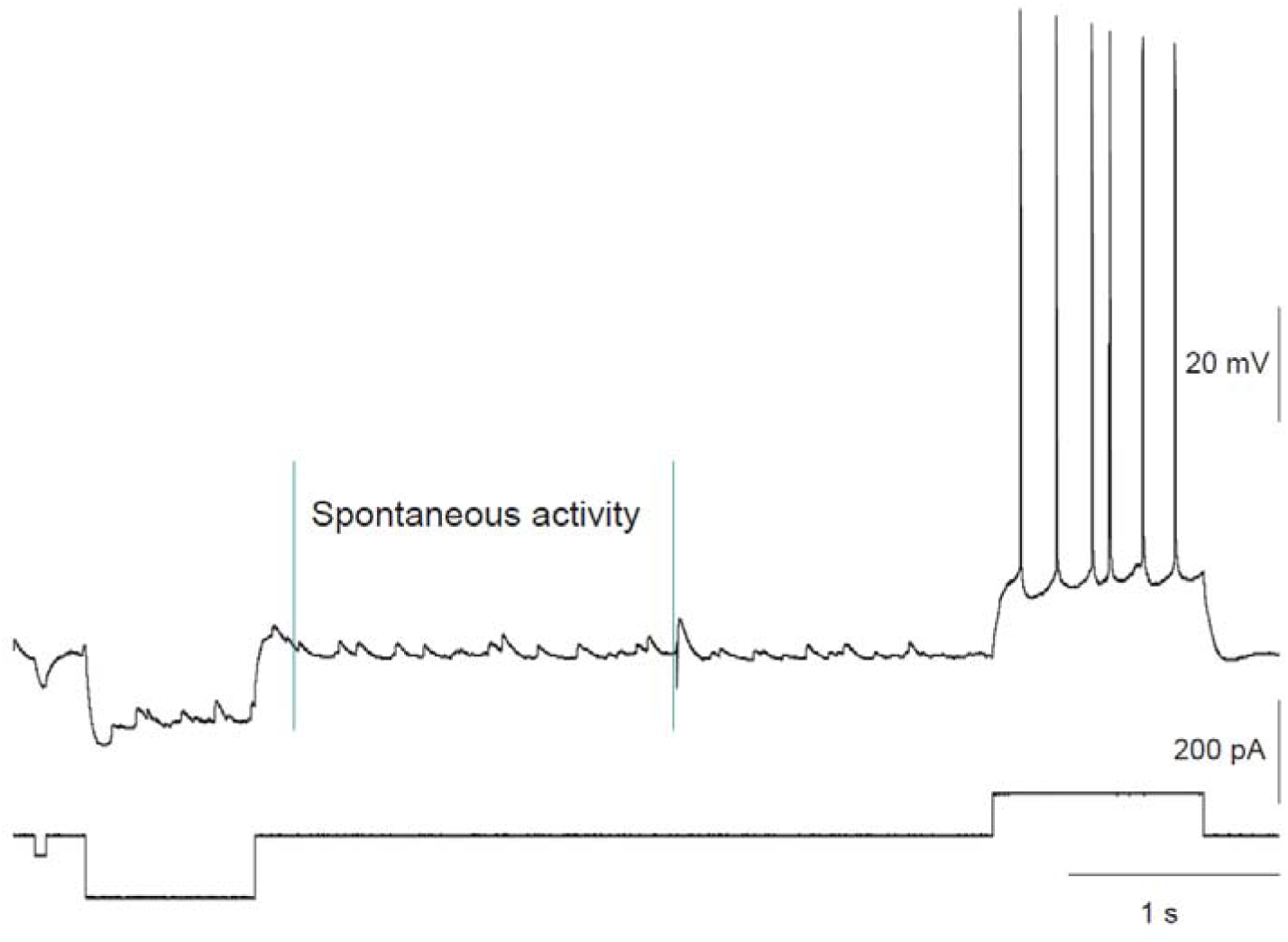
Measure of spontaneous activity. The period of spontaneous activity was measured after the large negative pulse of current and before the synaptic stimulation.

**Supplementary Figure 6.**
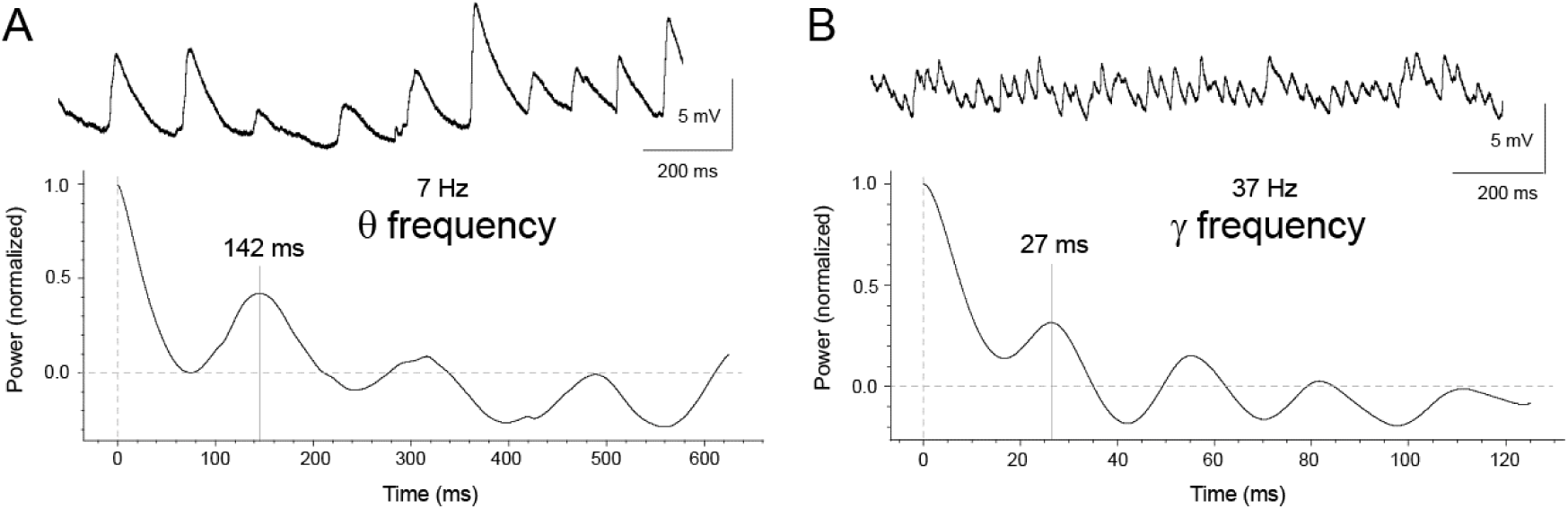
A. Spontaneous activity within the θ frequency range recorded in a CA1 pyramidal neuron. Top, representative trace. Bottom, auto-correlogram function of the voltage showing an oscillation over at least 3 cycles at 7 Hz. B. Spontaneous activity within the γ frequency range. Top, representative trace. Bottom, auto-correlogram function of the voltage showing an oscillation over at least 4 cycles at 37 Hz.

**Summary figure.**
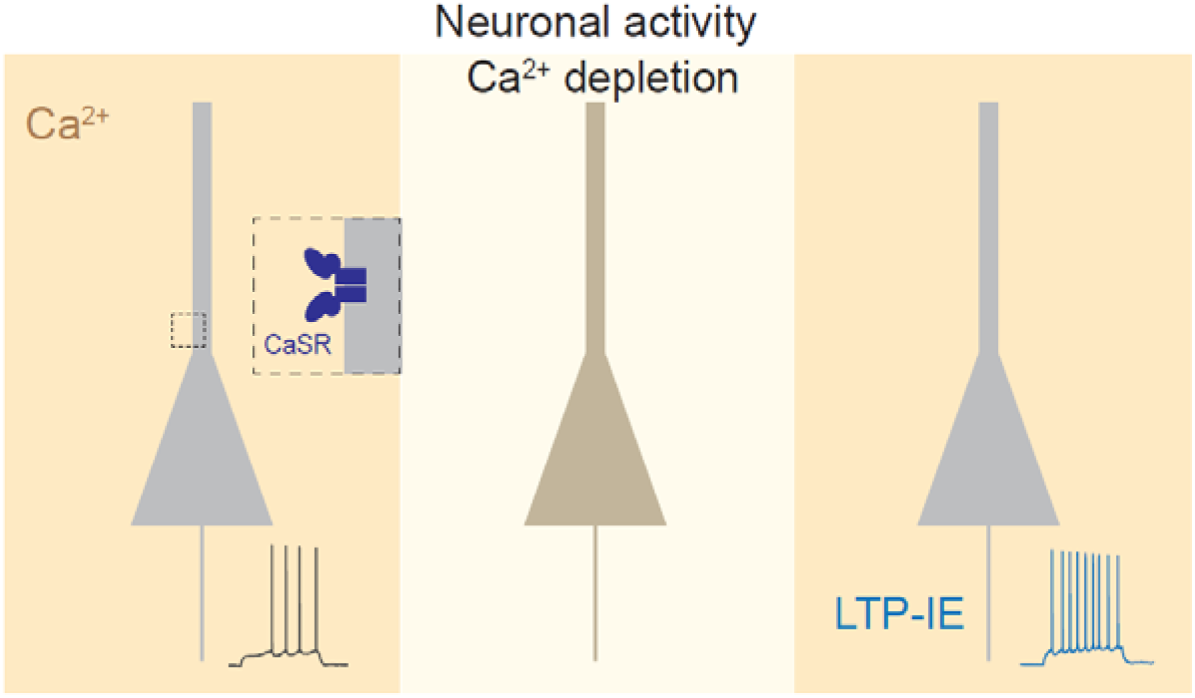

